# OPUS-Rota4: A Gradient-Based Protein Side-Chain Modeling Framework Assisted by Deep Learning-Based Predictors

**DOI:** 10.1101/2021.07.22.453446

**Authors:** Gang Xu, Qinghua Wang, Jianpeng Ma

**Affiliations:** Multiscale Research Institute of Complex Systems Fudan University Shanghai, 200433, China; Zhangjiang Fudan International Innovation Center Fudan University Shanghai, 201210, China; Shanghai AI Laboratory Shanghai, 200030, China; Verna and Marrs Mclean Department of Biochemistry and Molecular Biology Baylor College of Medicine Houston, Texas 77030, United States

**Keywords:** Protein Dihedral Angles Prediction, Protein Side-Chain Contact Map Prediction, Protein Side-Chain Modeling

## Abstract

Accurate protein side-chain modeling is crucial for protein folding and protein design. In the past decades, many successful methods have been proposed to address this issue. However, most of them depend on the discrete samples from the rotamer library, which may have limitations on their accuracies and usages. In this study, we report an open-source toolkit for protein side-chain modeling, named OPUS-Rota4. It consists of three modules: OPUS-RotaNN2, which predicts protein side-chain dihedral angles; OPUS-RotaCM, which measures the distance and orientation information between the side chain of different residue pairs; and OPUS-Fold2, which applies the constraints derived from the first two modules to guide side-chain modeling. In summary, OPUS-Rota4 adopts the dihedral angles predicted by OPUS-RotaNN2 as its initial states, and uses OPUS-Fold2 to refine the side-chain conformation with the constraints derived from OPUS-RotaCM. In this case, we convert the protein side-chain modeling problem into a side-chain contact map prediction problem. OPUS-Fold2 is written in Python and TensorFlow2.4, which is user-friendly to include other differentiable energy terms into its side-chain modeling procedure. In other words, OPUS-Rota4 provides a platform in which the protein side-chain conformation can be dynamically adjusted under the influence of other processes, such as protein-protein interaction. We apply OPUS-Rota4 on 15 FM predictions submitted by Alphafold2 on CASP14, the results show that the side chains modeled by OPUS-Rota4 are closer to their native counterparts than the side chains predicted by Alphafold2.

## Introduction

Protein side-chain modeling is an important task since the uniqueness of protein structure is largely determined by the packing of its side-chain conformation ^1^. In recent years, many successful programs have been proposed to address this issue ^1–11^.

The traditional protein side-chain modeling programs ^1, 4, 5, 10^ are composed of three key components: a rotamer library, an energy function, and a search method. One of the advantages of these methods is that they are very fast. Most of them can construct the side chains of a target within seconds. However, since these side-chain modeling methods depend on the discrete side-chain dihedral angles sampled from the rotamer library, the accuracy of the sampling candidates in the rotamer library determines the best performance these modeling methods can achieve.

With the development of deep learning techniques, some studies try to apply them to solve the protein side-chain modeling problem ^3, 11^. Our previous work OPUS-RotaNN in OPUS-Rota3 ^3^ tried to predict protein side-chain dihedral angles following the protocol we used to predict protein backbone torsion angles in OPUS-TASS ^12^. However, the accuracy of OPUS-RotaNN is worse than those of traditional side-chain modeling methods. We concluded that some new features may need to be designed to measure the local environment for each residue ^3^. Recently, DLPacker ^11^ used a 3DConv Neural Network to improve the accuracy of side-chain modeling by a large margin. Most importantly, the predicted density map from DLPacker is an excellent descriptor for the residue’s local environment measurement.

Protein structure prediction has become a hot topic since AlphaFold2 ^13^ from DeepMind achieved an astonishingly high performance in CASP14. Before that, the protein backbone structure prediction driven by contact map, which is used to describe if the Euclidean distance between two C_β_ atoms is less than 8.0 Å, is the most common way to deliver backbone conformation ^14^. Recently, trRosetta ^15^ supplemented the definition of contact map, including both distance and orientation information. The distance information is the traditional C_β_- C_β_ distance, and the orientation information between residues a and b includes 3 dihedrals (ω, θ_ab_, θ_ba_) and 2 angles (φ_ab_, φ_ba_). Here, ω represents the dihedral of C_αa_- C_βa_- C_βb_- C_αb_, θ_ab_ represents the dihedral of N_a_-C_αa_- C_βa_- C_βb_, φ_ab_ represents the angle of C_αa_- C_βa_- C_βb_. Since the backbone modeling driven by backbone contact map works well, we can develop the side-chain contact map for side-chain modeling accordingly. In this case, we convert the protein side-chain modeling problem from developing better scoring functions ^16^ to improving the accuracy of side-chain contact map prediction.

To generate protein 3D backbone structure from its corresponding backbone contact map, Crystallography and NMR System (CNS) ^17^ and pyRosetta ^18, 19^ are the most commonly used tools. In addition, the gradient-based backbone folding framework OPUS-Fold2 in our protein folding toolkit OPUS-X ^20^ can achieve comparable results to the Rosetta folding protocol in trRosetta ^15^. Moreover, OPUS-Fold2 is written in Python and TensorFlow2.4, which is user-friendly to any source-code level modification. Therefore, OPUS-Fold2 is suitable to be modified to deal with the side-chain modeling task.

In this paper, we propose an open-source toolkit for protein side-chain modeling, named OPUS-Rota4. It is comprised of three modules: OPUS-RotaNN2, OPUS-RotaCM, and OPUS-Fold2. OPUS-RotaNN2 includes some additional features, especially the local environment feature described by DLPacker ^11^, into its previous version OPUS-RotaNN ^3^, and delivers significantly better side-chain dihedral angles prediction than those from other state-of-the-art methods. OPUS-RotaCM reformats the input features from OPUS-RotaNN2 into 2D shape and uses them to predict the distance (C_β_- C_β_) and orientation (ω, θ_ab_, θ_ba_, φ_ab_, φ_ba_) information between the side chain of different residue pairs. OPUS-Fold2 used to be a gradient-based backbone folding framework ^20^, and it has been adjusted to deal with the side-chain modeling task in this work. It applies the constraints derived from the first two modules to guide side-chain modeling.

The contributions of this research can be summarized as follows:

1. The protein side-chain dihedral angles predicted by OPUS-RotaNN2 are significantly better than those predicted by other state-of-the-art methods in the literature, either measured by all residues or measured by core residues only.
2. We propose a side-chain contact map prediction method, OPUS-RotaCM, converting the protein side-chain modeling problem from developing better scoring functions to improving the accuracy of side-chain contact map prediction.
3. We develop a user-friendly gradient-based side-chain modeling framework, OPUS-Fold2, to refine the side-chain conformation. The protein side-chain conformation is adjustable when introducing the energy terms derived from other processes, and this may be useful for the corresponding process.
4. For non-native backbone side-chain modeling, OPUS-Rota4 can consistently deliver better results than other methods, showing its potential usage in structure prediction.

## Methods

### Framework of OPUS-Rota4

OPUS-Rota4 consists of three modules: OPUS-RotaNN2, OPUS-RotaCM and OPUS-Fold2. As shown in Figure 1, OPUS-RotaNN2 and OPUS-RotaCM share the same input features. 1D features are derived from protein backbone conformation, and there are 41 features in total. PC7 represents 7 physicochemical properties of each residue ^21^. PSP19 is derived from our previous work ^22, 23^, which classifies 20 residues into 19 rigid-body blocks. Here, PSP19 is a 19-d 0-1 vector, each dimension represents whether its corresponding rigid-body block is exists in the residue or not ^12^. SS3 and SS8 are two one-hot features that denote 3-state and 8-state secondary structure ^24^ of each residue, respectively. PP4 is the backbone torsion angles introduced as sin(*ϕ*), cos(*ϕ*), sin(*ψ*) and cos(*ψ*). 3DCNN is the side-chain density map predicted by DLPacker (OPUS) for each residue as its local environment descriptor. trRosetta100 is a 100-d feature which is used to describe backbone distance (C_β_- C_β_) and orientation (ω, θ_ab_, θ_ba_, φ_ab_, φ_ba_) contact information ^15^. CSF15 is the relative CSF position ^25^ of the backbone atoms of a specific residue at the local molecular coordinate system built in its contact counterpart ^3^.

**Figure 1.**
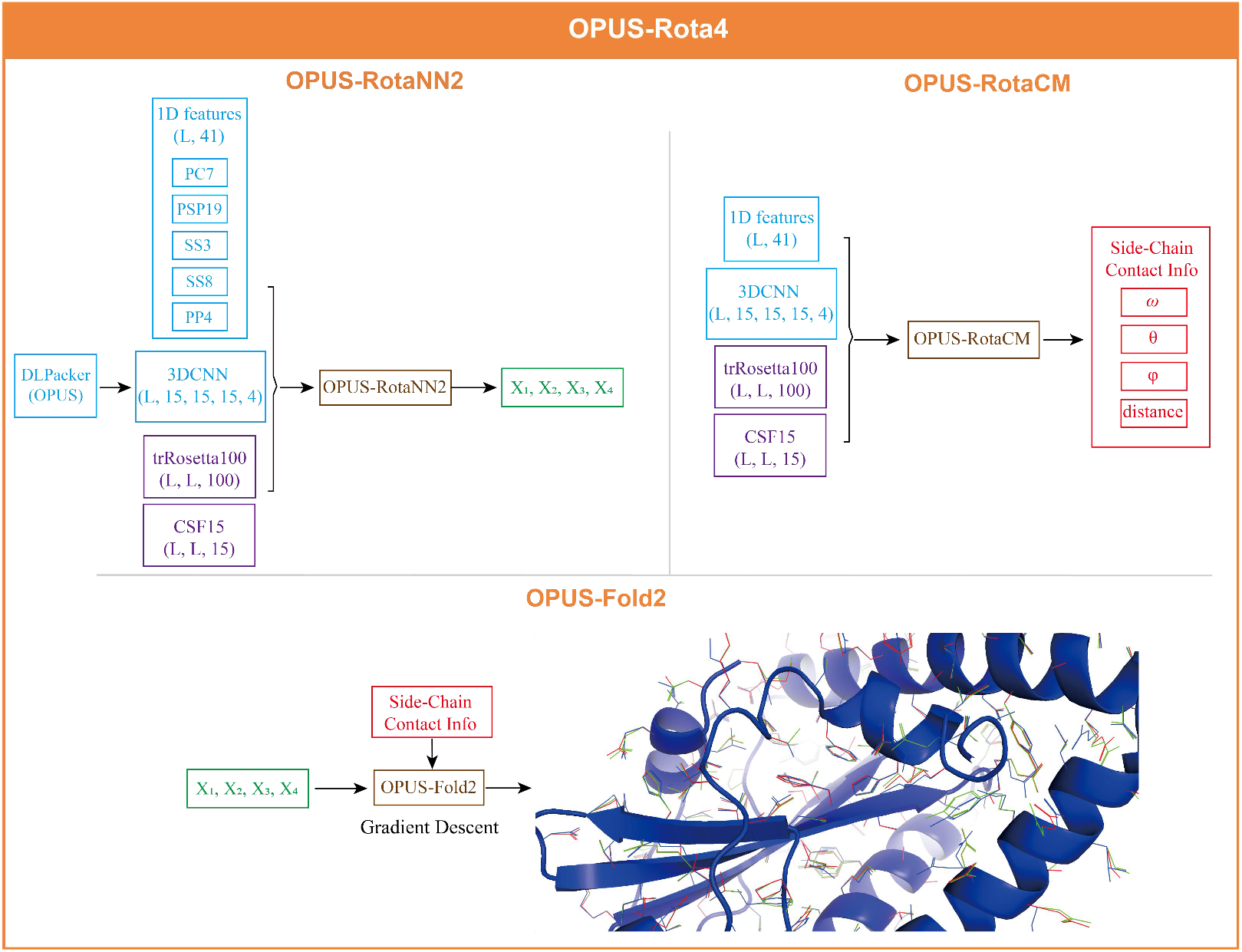
Flowchart of OPUS-Rota4. OPUS-Rota4 uses the dihedral angles predicted by OPUS-RotaNN2 as its initial states. Then, OPUS-Fold2 is used to refine the side-chain conformation with the side-chain contact constraints predicted by OPUS-RotaCM. *L* denotes the sequence length. The blue structure is the native state, the green structure is the initial state, and the red structure is the final prediction.

### DLPacker (OPUS)

Recently, a successful deep learning-based protein side-chain modeling method DLPacker ^11^ has been proposed, improving the accuracy of side-chain modeling by a large margin. Based on DLPacker, to capture the low-level feature more precisely, we respectively add 6 Residual Blocks to the two low-level feature pathways in the 3DConv U-Net ^26^ architecture of DLPacker, same as the DLPacker paper did in the high-level feature pathway. Meanwhile, we train 7 models and average their outputs to make the final prediction.

### OPUS-RotaNN2

The input features of OPUS-RotaNN2 can be categorized into 4 groups: 1D features, trRosetta100, CSF15 and 3DCNN. The output of OPUS-RotaNN2 contains 8 regression nodes: sin(*χ^1^*), cos(*χ^1^*), sin(*χ^2^*), cos(*χ^2^*), sin(*χ^3^*), cos(*χ^3^*), sin(*χ^4^*) and cos(*χ^4^*).

The neural network architecture of OPUS-RotaNN2 is shown in Figure 2, and it is mainly derived from the architecture of OPUS-TASS2 in OPUS-X paper ^20^. We use a stack of dilated residual-convolutional blocks to perform the feature extraction for 2D features, and use the attention mechanism ^27^ to sum up the multiply results of all residues with a specific residue and output a 128-d vector as its new feature. For 3DCNN, we use a MLP unit to generate a 512-d vector for each residue. Finally, we concatenate the three parts and feed them into the following modules which are identical to that in OPUS-TASS2 ^20^. We train 7 models and the median of their outputs is used to make the final prediction.

**Figure 2.**
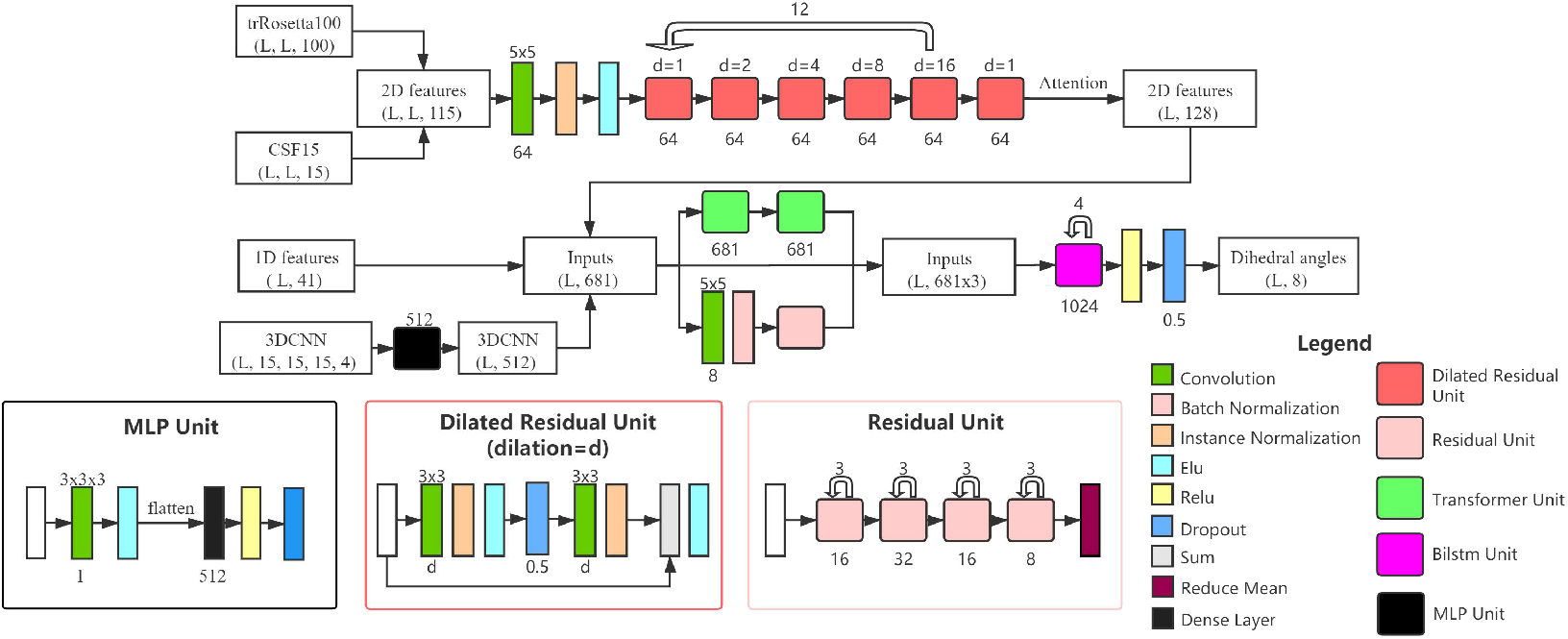
Framework of OPUS-RotaNN2. 2D features go through 61 dilated residual-convolutional blocks and an attention module ^27^, and output a 128-d vector for each residue. 3DCNN goes through a MPL unit and outputs a 512-d vector. Then, these two vectors and 1D features are concatenated to go through three modules: Resnet module ^28^, modified Transformer module ^27^ and bidirectional Long-Short-Term-Memory module ^29^. All strides in the residual units are set to be one. The batch size is also set to be one in OPUS-RotaNN2.

### OPUS-RotaCM

The input features of OPUS-RotaCM are identical to that of OPUS-RotaNN2. The output of OPUS-RotaCM is basically the same as that in trRosetta ^15^ but with some modifications. Instead of using C_α_ and C_β_ to measure the backbone conformation, we use the side-chain atoms as pseudo-C_α_ and C_β_ to measure the side-chain conformation. In general, for a specific residue, we use the side-chain atoms that are required for its side-chain dihedral angle *χ1* calculation. In this case, only *χ1* will be refined by OPUS-RotaCM. The detailed definitions of pseudo-C_α_ and C_β_ are shown in Supplementary Table S1. OPUS-RotaCM outputs one pseudo-C_β_- pseudo-C_β_ distance, 3 dihedrals (ω, θ_ab_, θ_ba_) and 2 angles (φ_ab_, φ_ba_) between residues a and b. The distance ranges from 2 to 20 Å, segmented into 36 bins with 0.5 Å interval, and with one extra bin represents the >20 Å case. ω and θ range from −180 to 180°, segmented into 24 bins with 15° interval, and with one extra bin represents the non-contact case. φ ranges from 0 to 180°, segmented into 12 bins with 15° interval, and with one extra bin represents the non-contact case.

The neural network architecture of OPUS-RotaCM is shown in Figure 3. We train 7 models and the average of their outputs is used to make the final prediction.

**Figure 3.**
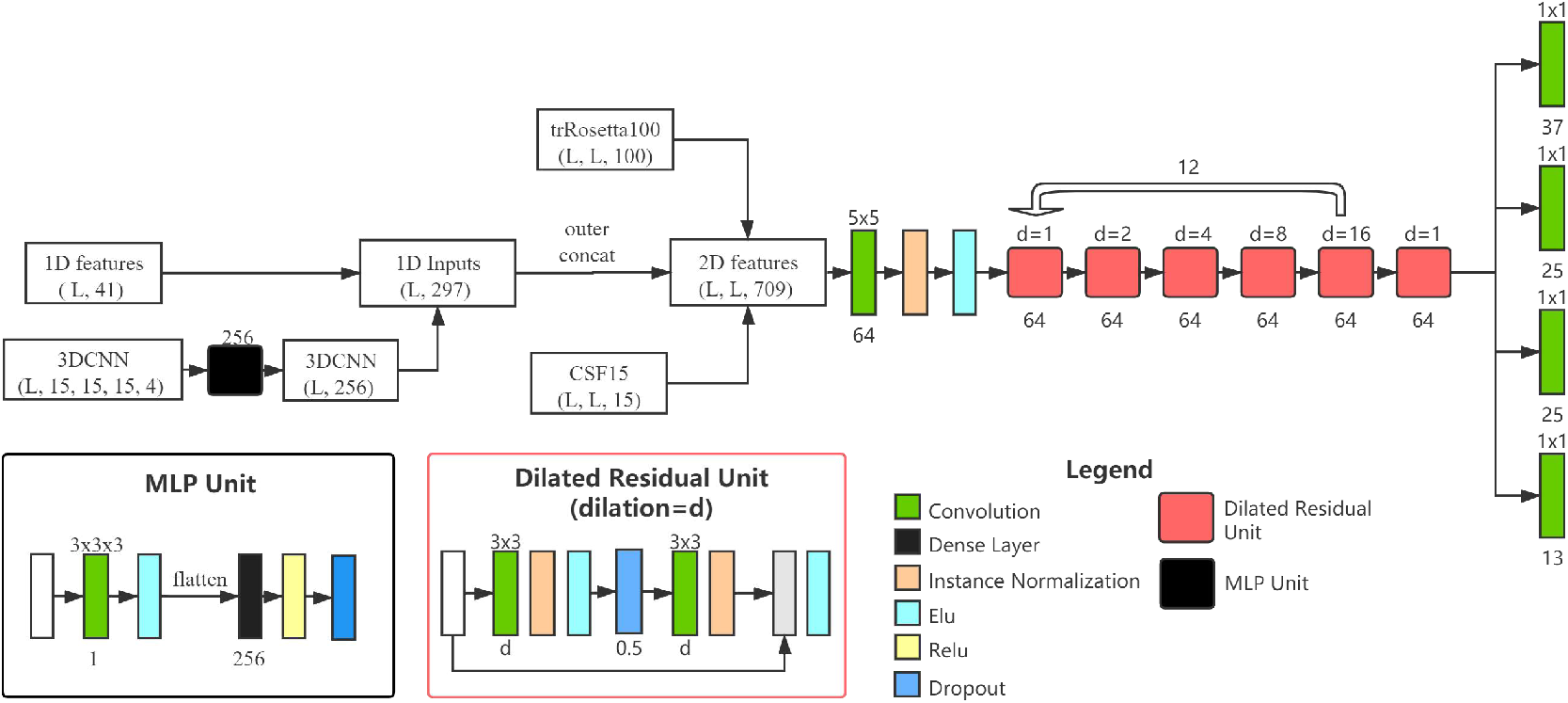
Framework of OPUS-RotaCM. 3DCNN goes through a MPL unit and outputs a 256-d vector. Then 1D features and 3DCNN are concatenated to form a 297-d 1D inputs vector. We use outer concatenation function to convert 1D inputs (L, 297) into 2D features (L, L, 594). After concatenating with trRosetta100 and CSF15, they go through a stack of 61 dilated residual-convolutional blocks. The batch size is set to be one in OPUS-RotaCM.

### OPUS-Fold2

OPUS-Fold2 used to be a gradient-based backbone folding framework ^20^ and it has been modified to be a gradient-based side-chain modeling framework in this research. The variables of OPUS-Fold2 are the side-chain dihedral angles (*χ^1^*, *χ^2^*, *χ^3^* and *χ^4^*) of all residues. OPUS-Fold2 optimizes its variables to minimize the loss function derived from the output of OPUS-RotaCM.

The predictions from OPUS-RotaNN2 are set to be the initial states of *χ^1^*, *χ^2^*, *χ^3^* and *χ^4^*. Same as the backbone modeling version in OPUS-X ^20^, the loss function of OPUS-Fold2 in this research is defined as follows:

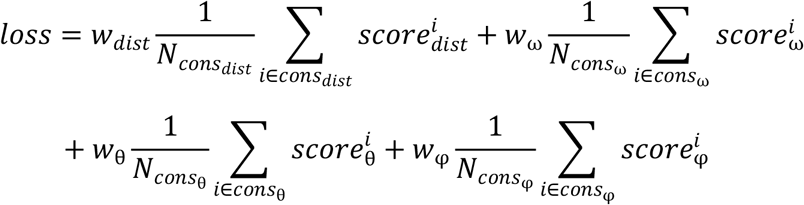

*cons_dist_* is the collection of distance constraints with probability *P_4≤dist<20_ ≥ 0.05*. *cons_ω_* and *cons_θ_* are the collections of ω and θ constraints with probability *P_contact_ ≥ 0.55*. *cons_φ_* is the collections of φ constraints with probability *P_contact_ ≥ 0.65*. *w_dist_*, *w_ω_*, *w_θ_* and *w_φ_* are the weight of each term, which are set to be 5, 4, 4 and 4, respectively.

For distance distribution, we use the following equation: 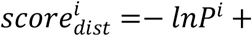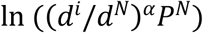, where *P^i^* is the probability of the *i*th bin, *d^i^* is the distance of the *i*th distance bin, α is 1.57 ^30^ and *N* is the bin [19.5, 20]. For orientation distribution, we use the following equation: 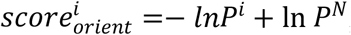, where *N* is bin [165°, 180°]. Then, cubic spline curve is generated to make each distribution differentiable.

OPUS-Fold2 is implemented based on TensorFlow2.4 ^31^. Adam ^32^ optimizer is used to optimize the loss function with an initial learning rate of 1.0, 500 epochs are performed. The side-chain conformation with the lowest loss during the optimization is considered as the final prediction.

### Datasets

OPUS-RotaNN2 and OPUS-RotaCM use the same training and validation sets as those in OPUS-RotaNN ^3^, which were culled from the PISCES server ^33^ by SPOT-1D ^21^ on February 2017 with following constraints: R-free <1, resolution > 2.5 Å, and sequence identity < 25%. There are 10024 proteins in the training set and 983 proteins in the validation set.

To evaluate the side-chain modeling for native backbone structure, we use three independent test sets. CASPFM (56), collected by SAINT ^34^, contains 56 Template-Free Modelling (FM) targets obtained from CASP10 to CASP13. CASP14 (15), collected by OPUS-X ^20^, contains 15 FM targets downloaded from the CASP website (http://predictioncenter.org). CAMEO (60), collected by OPUS-Rota3 ^3^, contains 60 hard targets (we discard one target with over 900 residues in length) released between January 2020 and July 2020 from the CAMEO website ^35^. To evaluate the side-chain modeling for non-native backbone structure, we collect the predictions submitted by Alphafold2 ^13^ for the targets in CASP14 (15). This non-native backbone dataset is denoted as CASP14-AF2 (15). The average TM-score^36^ of these 15 predictions from their native counterparts is 0.85.

### Performance Metrics

MAE (*χ^1^*), MAE (*χ^2^*), MAE (*χ^3^*) and MAE (*χ^4^*) are used to measure the mean absolute error (MAE) of *χ^1^*, *χ^2^*, *χ^3^* and *χ^4^* between the native value and the predicted one, respectively. ACC is used to measure the percentage of correct prediction with a tolerance criterion 20° for all side-chain dihedral angles (from *χ^1^* to *χ^4^*). RMSD is calculated by the *Superimposer* function in Biopython ^37^ residue-wisely. Paired t-test is used to get the significance value P for the residue-wise comparison. Following FASPR ^10^, the residue with more than 20 residues, between which the C_β_-C_β_ distance is within 10 Å, is defined as core residue. The C_α_ atom is used for Gly. In summary, 25% residues in CAMEO (60), 18% residues in CASPFM (56) and 17% residues in CASP14 (15) are defined as core residue.

## Results

### Input Feature Study

To evaluate the importance of 4 input feature groups in OPUS-RotaNN2, we add them to the input of OPUS-RotaNN2 and train the model one by one. As the results shown in Table 1, in terms of MAE (*χ1*), MAE (*χ2*), MAE (*χ3*), MAE (*χ4*) and ACC, the accuracy of OPUS-RotaNN2 is gradually increased after introducing these feature groups into its input. OPUS-RotaNN2 finally achieves the best performance when using 4 input feature groups together.

**Table 1.** The performance of OPUS-RotaNN2 after introducing corresponding input feature groups.

| | MAE ($\chi_1$) | MAE ($\chi_2$) | MAE ($\chi_3$) | MAE ($\chi_4$) | ACC |
| --- | --- | --- | --- | --- | --- |
| CAMEO (60) |  |  |  |  |  |
| 1D features | 35.51 | 43.28 | 57.84 | 51.37 | 36.53% |
| +trRosetta100 | 31.52 | 41.69 | 57.07 | 51.49 | 40.32% |
| +CSF15 | 30.25 | 41.31 | 57.50 | 51.57 | 40.50% |
| +3DCNN | **21.77** | **31.24** | **49.64** | **47.66** | **55.63%** |
| CASPFM (56) |  |  |  |  |  |
| 1D features | 31.19 | 39.94 | 53.02 | 49.01 | 41.93% |
| +trRosetta100 | 27.15 | 37.44 | 53.09 | 49.22 | 46.42% |
| +CSF15 | 25.75 | 36.25 | 53.17 | 49.63 | 46.94% |
| +3DCNN | **18.85** | **28.80** | **44.89** | **45.06** | **57.97%** |
| CASP14 (15) |  |  |  |  |  |
| 1D features | 42.04 | 46.35 | 54.31 | 39.78 | 28.56% |
| +trRosetta100 | 37.73 | 45.67 | 54.41 | **39.19** | 30.42% |
| +CSF15 | 35.56 | 45.20 | 53.07 | 39.46 | 31.25% |
| +3DCNN | **28.35** | **40.34** | **51.41** | 41.25 | **41.25%** |

### Performance of Different Side-Chain Modeling Methods

We compare the direct prediction results from OPUS-RotaNN2 and the final refined results from OPUS-Rota4 with those from three rotamer library-based methods FASPR ^10^, SCWRL4 ^5^ and OSCAR-star ^4^, and two deep learning-based methods OPUS-RotaNN ^3^ and DLPacker ^11^. In terms of MAE (*χ1*), MAE (*χ2*), MAE (*χ3*), MAE (*χ4*) and ACC, OPUS-RotaNN2 and OPUS-Rota4 outperform other methods by a large margin, either measured by all residues (Table 2) or measured by core residues only (Supplementary Table S2). Note that, the difference between OPUS-RotaNN2 and OPUS-Rota4 only exists in *χ1* since OPUS-Fold2 uses the side-chain contact constraints derived from the pseudo-C_α_ and C_β_ that are required for *χ1* calculation from OPUS-RotaCM. As shown in Table 3, in terms of residue-wise RMSD, OPUS-Rota4 also delivers better results than other methods for all residues and core residues.

**Table 2.** The performance of different side-chain modeling methods on three native backbone test sets measured by all residues.

| | MAE ($\chi_1$) | MAE ($\chi_2$) | MAE ($\chi_3$) | MAE ($\chi_4$) | ACC |
| --- | --- | --- | --- | --- | --- |
| CAMEO (60) |  |  |  |  |  |
| FASPR | 29.15 | 42.36 | 57.01 | 57.93 | 49.10% |
| SCWRL4 | 29.01 | 42.88 | 57.25 | 57.17 | 49.48% |
| OSCAR-star | 27.29 | 41.97 | 56.08 | 57.66 | 49.91% |
| OPUS-RotaNN | 33.28 | 42.47 | 57.68 | 51.39 | 37.83% |
| DLPacker | 24.11 | 39.60 | 63.84 | 68.10 | 52.19% |
| OPUS-RotaNN2 | 21.61 | 31.13 | 49.79 | 47.78 | 55.61% |
| OPUS-Rota4 | **21.34** | **31.13** | **49.79** | **47.78** | **57.35%** |
| CASPFM (56) |  |  |  |  |  |
| FASPR | 26.63 | 39.75 | 53.40 | 54.81 | 53.11% |
| SCWRL4 | 27.09 | 40.44 | 52.67 | 54.61 | 53.17% |
| OSCAR-star | 24.53 | 37.43 | 50.51 | 52.99 | 54.92% |
| OPUS-RotaNN | 29.41 | 38.93 | 53.33 | 49.19 | 42.86% |
| DLPacker | 21.35 | 37.79 | 61.05 | 66.78 | 55.26% |
| OPUS-RotaNN2 | 18.85 | 28.50 | 44.88 | 44.87 | 58.17% |
| OPUS-Rota4 | **18.46** | **28.50** | **44.88** | **44.87** | **60.42%** |
| CASP14 (15) |  |  |  |  |  |
| FASPR | 35.80 | 48.72 | 56.59 | 45.19 | 36.34% |
| SCWRL4 | 35.27 | 48.13 | 58.37 | 48.15 | 36.57% |
| OSCAR-star | 34.45 | 48.10 | 56.70 | 42.28 | 36.76% |
| OPUS-RotaNN | 39.57 | 45.67 | 53.80 | **39.77** | 27.31% |
| DLPacker | 30.99 | 48.21 | 65.14 | 70.83 | 40.05% |
| OPUS-RotaNN2 | **28.21** | 40.14 | 51.93 | 40.76 | 41.16% |
| OPUS-Rota4 | 28.33 | **40.14** | **51.93** | 40.76 | **43.38%** |

**Table 3.**
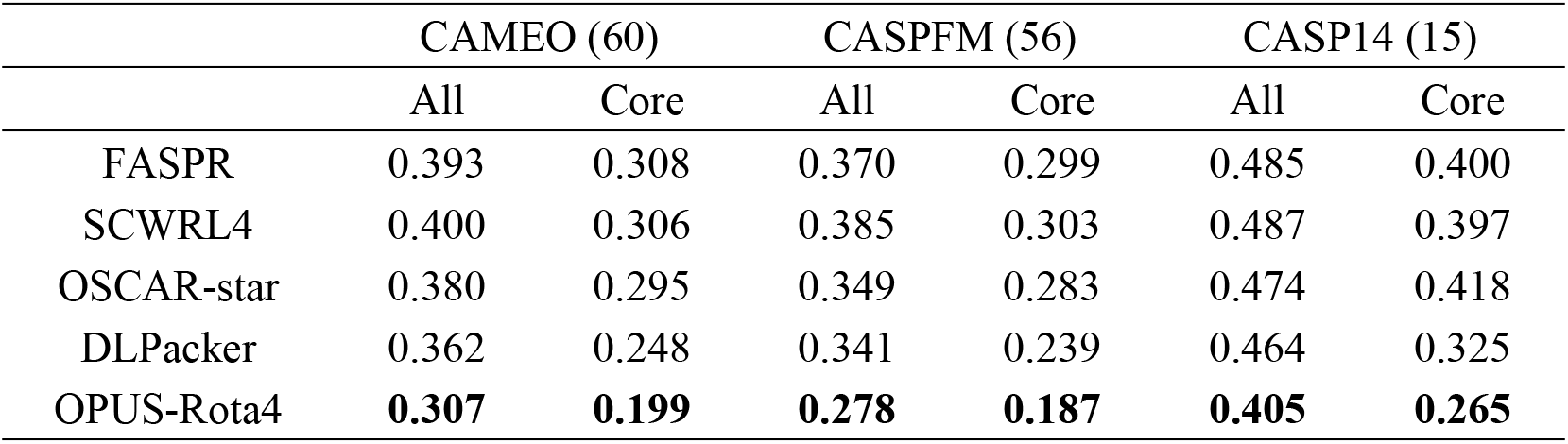
The RMSD results of different side-chain modeling methods on three native backbone test sets.

### Side-Chain Modeling for Non-Native Backbone Structure

We evaluate the performance of different side-chain modeling methods on CASP14-AF2 (15). The MAE (*χ1*), MAE (*χ2*), MAE (*χ3*), MAE (*χ4*) and ACC results are shown in Table 4. In terms of ACC, OPUS-Rota4 outperforms other methods, including the original side chains submitted by Alphafold2 ^13^. The RMSD result of each method and its significance value comparing with the result from OPUS-Rota4 is listed in Table 5. The results show that OPUS-Rota4 significantly outperforms other methods on all residues and core residues. The detailed comparisons between OPUS-Rota4 and Alphafold2 for each target are listed in Supplementary Table S3. In terms of RMSD results for all residues, the side chains modeled by OPUS-Rota4 are closer to their native counterparts than the original side chains of Alphafold2’s predictions on 13 out of 15 targets in CASP14-AF2 (15).

**Table 4.** The performance of different side-chain modeling methods on CASP14-AF2 (15).

| | MAE ($\chi_1$) | MAE ($\chi_2$) | MAE ($\chi_3$) | MAE ($\chi_4$) | ACC |
| --- | --- | --- | --- | --- | --- |
| All |  |  |  |  |  |
| Alphafold2 | **40.14** | 57.46 | 71.25 | 41.79 | 30.56% |
| FASPR | 44.73 | 52.92 | 57.91 | 46.12 | 29.91% |
| SCWRL4 | 45.51 | 51.86 | 57.43 | 48.86 | 29.91% |
| OSCAR-star | 43.83 | 52.13 | 58.14 | 48.46 | 30.09% |
| DLPacker | 43.04 | 54.90 | 67.63 | 70.41 | 30.05% |
| OPUS-RotaNN2 | 42.00 | 46.48 | 55.81 | 39.41 | 30.88% |
| OPUS-Rota4 | 42.06 | **46.48** | **55.81** | **39.41** | **32.13%** |
| Core |  |  |  |  |  |
| AlphaFold2 | **26.93** | 53.46 | 61.61 | **32.26** | 47.51% |
| FASPR | 30.80 | 53.59 | 54.12 | 59.18 | 43.09% |
| SCWRL4 | 31.13 | 51.25 | 54.48 | 50.34 | 44.20% |
| OSCAR-star | 32.26 | 51.25 | 51.55 | 63.89 | 43.65% |
| DLPacker | 30.55 | 49.02 | 60.24 | 51.03 | 45.03% |
| OPUS-RotaNN2 | 28.78 | 40.87 | 59.03 | 34.07 | 50.28% |
| OPUS-Rota4 | 28.51 | **40.87** | **59.03** | 34.07 | **51.38%** |

**Table 5.** The RMSD results of different side-chain modeling methods on CASP14-AF2 (15).

|  | All |  | Core |  |
| --- | --- | --- | --- | --- |
|  | RMSD | P<sub>RMSD</sub> | RMSD | P<sub>RMSD</sub> |
| AlphaFold2 | 0.588 | 1.3E-12 | 0.472 | 5.5E-04 |
| FASPR | 0.574 | 2.5E-09 | 0.484 | 1.4E-05 |
| SCWRL4 | 0.585 | 3.9E-14 | 0.489 | 1.0E-05 |
| OSCAR-star | 0.569 | 5.9E-08 | 0.483 | 3.0E-05 |
| DLPacker | 0.576 | 1.1E-13 | 0.449 | 5.9E-04 |
| OPUS-Rota4 | **0.535** | - | **0.407** | - |

### Performance of OPUS-Fold2 on Side-Chain Modeling

OPUS-Fold2 is a gradient-based side-chain modeling framework, and it is able to refine all side-chain atoms. In OPUS-Rota4, we use the prediction from OPUS-RotaCM to refine the side-chain dihedral *χ^1^* only. For *χ^2^* refinement, we set the pseudo-C_α_ and C_β_ to be those atoms required for *χ^2^* calculation and retrain the OPUS-RotaCM model. Then we use the predicted *χ^2^* constraints from the modified OPUS-RotaCM to further refine the prediction from OPUS-Rota4, the results in Supplementary Table S4 show that the *χ^2^* constraints are not accurate enough to improve the *χ^2^* accuracy. It indicates that the side-chain contact map prediction for more flexible dihedral *χ^2^* is more challenging.

To verify the side-chain modeling ability of OPUS-Fold2, we use the side-chain contact map constraints derived from the native structures for *χ^1^* to *χ^4^* to guide side-chain modeling. As shown in Table 6, with the correct constraints, OPUS-Fold2 can guide the side chains to their proper places.

**Table 6.**
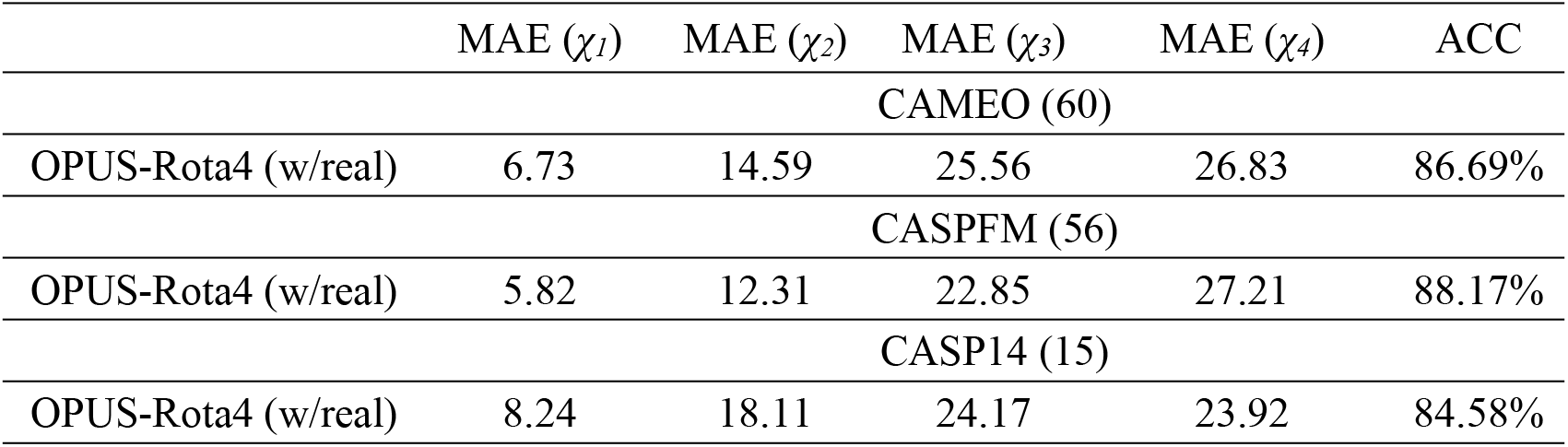
The performance of OPUS-Fold2 using the side-chain contact map constraints derived from the native structures.

### Case study

We show some successful and failed cases of OPUS-Rota4 side-chain modeling results in Figure 4 and Figure 5, respectively. It shows that side-chain modeling for the longish loop area may need to be further refined.

**Figure 4.**
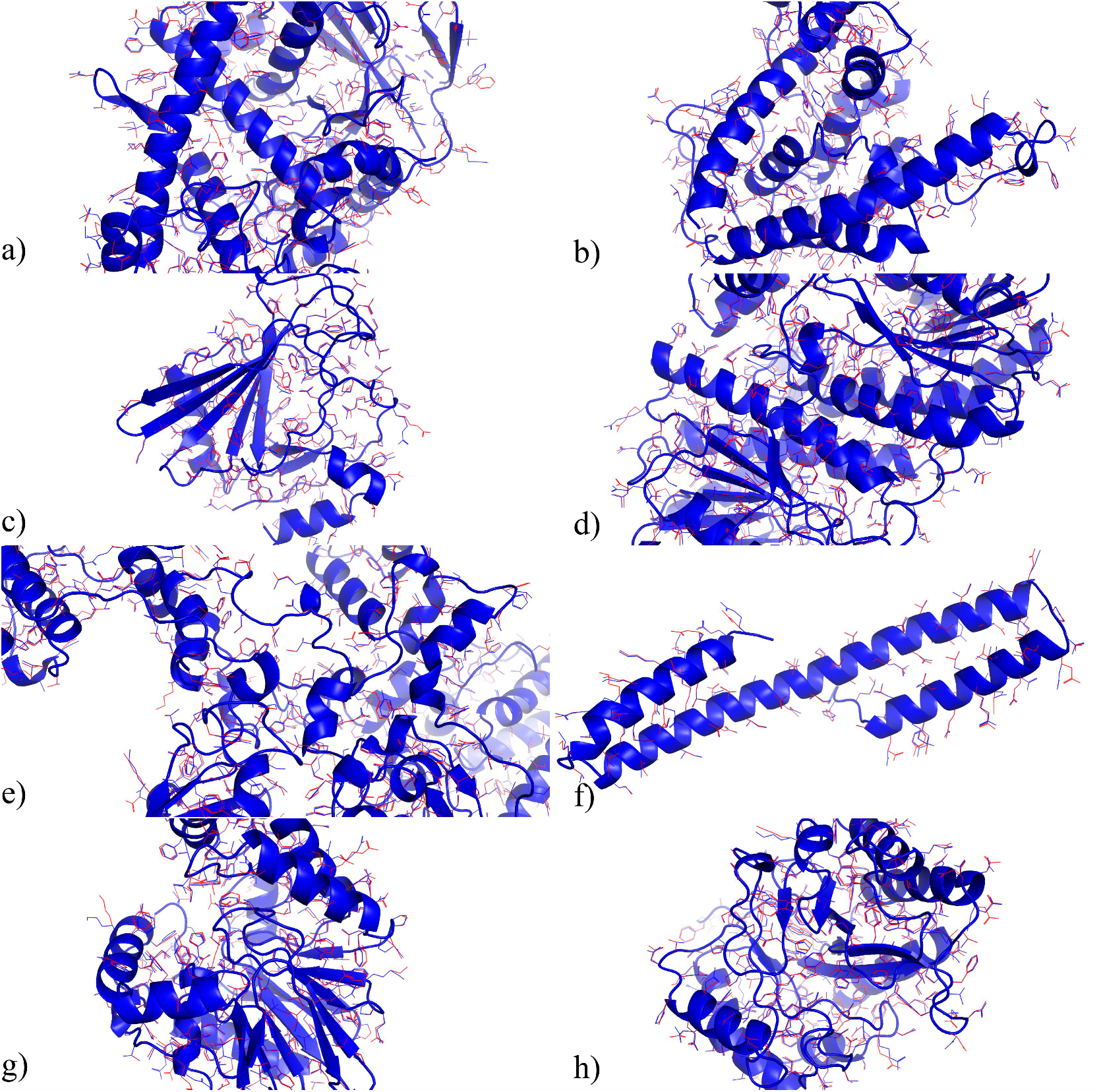
Successful side-chain modeling examples of OPUS-Rota4. a) T1037-D1 (Length: 404, RMSD: 0.418), b) T1041-D1 (Length: 241, RMSD: 0.421), c) T1090-D1 (Length: 177, RMSD: 0.214), d) 2020-03-14_00000031_1 (Length: 545, RMSD: 0.24), e) 2020-03-21_00000182_1 (Length: 658, RMSD: 0.227), f) 2020-04-18_00000132_1 (Length: 98, RMSD: 0.285), g) 2020-05-09_00000226_1 (Length: 300, RMSD: 0.191), h) 2020-05-16_00000125_1 (Length: 257, RMSD: 0.188). The blue structure is the native state and the red structure is the prediction.

**Figure 5.**
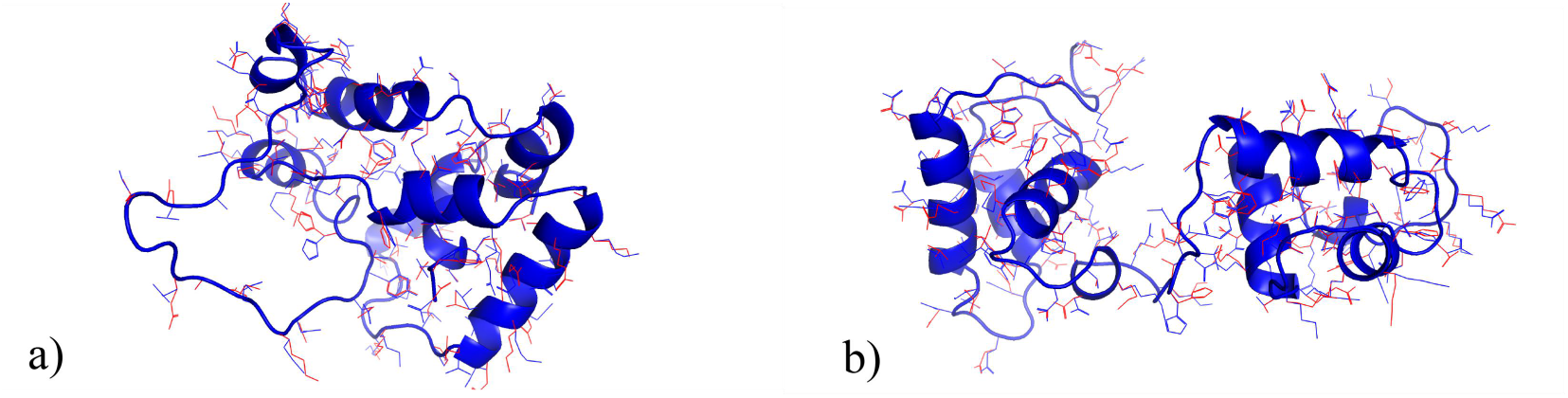
Failed side-chain modeling examples of OPUS-Rota4. a) T1027-D1 (Length: 127, RMSD: 0.521), b) 2020-06-27_00000154_1 (Length: 141, RMSD: 0. 632). The blue structure is the native state and the red structure is the prediction.

## Concluding Discussion

Protein side-chain modeling is a crucial task since many important biological processes depend on the interaction of protein side chains. In this paper, we develop an open-source toolkit for protein side-chain modeling, named OPUS-Rota4. It includes a side-chain dihedral angles predictor, namely OPUS-RotaNN2; a side-chain contact map predictor, namely OPUS-RotaCM; and a gradient-based side-chain modeling framework, namely OPUS-Fold2.

The performance of traditional rotamer library-based side-chain modeling methods are limited by the accuracy of the discrete sampling candidates in the rotamer library and the precision of their scoring functions, therefore, there may be an upper bound for these methods. As shown in Table 2, Table 3, Table 4 and Table 5, FASPR ^10^ and SCWRL4 ^5^ achieve similar performance on both native backbone and non-native backbone test sets. OSCAR-star ^4^ is better than those two methods for using more effective scoring function. One of the advantages of these methods is that they can deliver the results within seconds, which is suitable for iterative side-chain construction.

For deep learning-based side-chain modeling method, the major issue is how to define the local environment for each residue properly ^3^. Benefit from DLPacker ^11^, a recently proposed method which uses a 3DConv Neural Network to output the side-chain density map for each residue, we use the side-chain density map as the local environment descriptor in OPUS-RotaNN2 and significantly improve the final prediction (Table 1).

As shown in Table 2, Supplementary Table S2 and Table 3, the performance of OPUS-RotaNN2 is significantly better than the performance of FASPR ^10^, SCWRL4 ^5^, OSCAR-star ^4^, OPUS-RotaNN ^3^ and DLPacker ^11^ on three native backbone test sets, either measured by all residues or measured by core residues only. For non-native backbone side-chain modeling, which is especially useful in protein structure prediction, OPUS-RotaNN2 and OPUS-Rota4 also achieve better results than these methods (Table 4 and Table 5). In addition, comparing with the original side chains submitted by Alphafold2 ^13^, the side chains modeled by OPUS-Rota4 are closer to their native counterparts on 13 out of 15 targets in CASP14-AF2 (15) (Supplementary Table S3). We believe that the side chains that are closer to their native states may give a positive feedback to refine their corresponding backbones further.

Predicting accurate protein side-chain dihedral angles directly is important, but what is more crucial is how to refine them in a differentiable manner. On the one hand, the accuracy of side-chain dihedral angles can be further improved by other differentiable energy terms. On the other hand, making side chains adjustable may be benefit for some other processes that can be introduced into the energy function, such as protein-protein interaction.

Inspired by the successful usage of backbone contact map in protein backbone structure prediction ^14, 15^, we develop a side-chain contact map predictor OPUS-RotaCM. From another point of view, side-chain contact map that includes distance and orientation information can be considered as a more accurate scoring function. In OPUS-RotaCM, we use the side-chain atoms that are required for the side-chain dihedral angle *χ^1^* calculation as pseudo-C_α_ and C_β_ to measure the side-chain conformation. As shown in Table 2, Supplementary Table S2 and Table 4, the *χ^1^* accuracy of OPUS-RotaNN2 can be further refined by the constraints derived from OPUS-RotaCM (OPUS-RotaNN2 versus OPUS-Rota4). However, for *χ^2^* refinement, the predicted side-chain contact map constraints obtained by following the same training and inference protocol for *χ1* are not accurate enough (Supplementary Table S4), which means *χ^2^* is more flexible than *χ^1^*, and more new features may need to be introduced.

OPUS-Fold2 used to be a gradient-based framework for backbone folding ^20^, and it has been modified to be a side-chain modeling framework in this paper. As shown in Table 6, OPUS-Fold2 can guide the side chains to their proper places with the correct side-chain contact constraints, showing the effectiveness of its side-chain modeling ability. In this case, we can improve the protein side-chain modeling accuracy by improving the accuracy of side-chain contact map prediction other than developing better scoring functions ^16^.

## Availability and implementation

The code and pre-trained models of OPUS-Rota4 can be downloaded from https://github.com/OPUS-MaLab/opus_rota4.

## Acknowledgements

The work was partially supported by Shanghai Municipal Science and Technology Major Project (No.2018SHZDZX01), and ZJLab. QW thanks the Welch Foundation (Q-1826) for support. JM thanks the support from the Welch Foundation (Q-1512).

## Author contributions

Gang Xu and Jinapeng Ma designed the project. Gang Xu conducted the experiments. All authors contributed to the manuscript composing.

