## Supplementary material for "OPUS-Rota4: A Gradient-Based Protein Side-Chain Modeling Framework Assisted by Deep Learning-Based Predictors": SI

**Table S1. Definitions of pseudo-$C_\alpha$ and $C_\beta$ in OPUS-RotaCM.**

| Residue type | pseudo-$C_\alpha$ | pseudo- $C_\beta$ |
|---|---|---|
| G | $C_\alpha$ | $C_\beta$ (Constructed as Ala) |
| A | $C_\alpha$ | $C_\beta$ |
| S | $C_\beta$ | OG |
| C | $C_\beta$ | SG |
| V | $C_\beta$ | CG1 |
| I | $C_\beta$ | CG1 |
| L | $C_\beta$ | CG |
| T | $C_\beta$ | OG1 |
| R | $C_\beta$ | CG |
| K | $C_\beta$ | CG |
| D | $C_\beta$ | CG |
| E | $C_\beta$ | CG |
| N | $C_\beta$ | CG |
| Q | $C_\beta$ | CG |
| M | $C_\beta$ | CG |
| H | $C_\beta$ | CG |
| P | $C_\alpha$ | CG |
| F | $C_\beta$ | CG |
| Y | $C_\beta$ | CG |
| W | $C_\beta$ | CG |

**Table S2. The performance of different side-chain modeling methods on three native backbone test sets measured by core residues.**

| | MAE ($\chi_1$) | MAE ($\chi_2$) | MAE ($\chi_3$) | MAE ($\chi_4$) | ACC |
|---|---|---|---|---|---|
| | | CAMEO (60) | | | |
| FASPR | 17.24 | 38.16 | 53.07 | 57.10 | 63.66% |
| SCWRL4 | 16.95 | 37.98 | 49.80 | 49.09 | 65.85% |
| OSCAR-star | 16.31 | 37.71 | 49.01 | 60.44 | 64.30% |
| OPUS-RotaNN | 26.67 | 43.19 | 62.42 | 44.96 | 47.46% |
| DLPacker | 12.45 | 31.14 | 52.49 | 59.97 | 70.73% |
| OPUS-RotaNN2 | **10.64** | 22.94 | 40.63 | 39.38 | 75.85% |
| OPUS-Rota4 | 10.71 | **22.94** | **40.63** | **39.38** | **76.61%** |
| | | CASPFM (56) | | | |
| FASPR | 14.77 | 36.43 | 49.95 | 72.28 | 67.16% |
| SCWRL4 | 15.19 | 34.51 | 45.63 | 61.82 | 68.30% |
| OSCAR-star | 14.78 | 33.09 | 41.18 | 63.54 | 68.46% |
| OPUS-RotaNN | 24.07 | 38.24 | 58.02 | 52.20 | 52.00% |
| DLPacker | 11.45 | 27.19 | 48.45 | 54.94 | 71.48% |
| OPUS-RotaNN2 | **9.31** | 20.51 | 31.08 | 41.94 | 77.18% |
| OPUS-Rota4 | 9.37 | **20.51** | **31.08** | **41.94** | **78.16%** |
| | | CASP14 (15) | | | |
| FASPR | 22.52 | 47.08 | 53.02 | 46.34 | 50.83% |
| SCWRL4 | 21.45 | 47.74 | 46.80 | 34.68 | 51.66% |
| OSCAR-star | 25.28 | 50.40 | 56.02 | 30.42 | 49.45% |
| OPUS-RotaNN | 32.51 | 48.18 | 50.18 | 33.62 | 34.25% |
| DLPacker | 17.66 | 39.78 | 69.73 | 42.77 | 60.50% |
| OPUS-RotaNN2 | 13.44 | 32.48 | 39.67 | 27.91 | 64.92% |
| OPUS-Rota4 | **13.16** | **32.48** | **39.67** | **27.91** | **66.30%** |

**Table S3. The RMSD results of OPUS-Rota4 and Alphafold2 on CASP14-AF2 (15).**

| | | | OPUS-Rota4 | Alphafold2 | OPUS-Rota4 | Alphafold2 |
|---|---|---|---|---|---|---|
| | TM-score | Length | All | | Core | |
| T1090-D1 | 0.96 | 177 | **0.346** | 0.409 | **0.244** | 0.351 |
| T1037-D1 | 0.96 | 404 | **0.542** | 0.590 | **0.404** | 0.486 |
| T1041-D1 | 0.95 | 241 | **0.529** | 0.607 | **0.438** | 0.497 |
| T1042-D1 | 0.94 | 276 | **0.550** | 0.630 | **0.487** | 0.560 |
| T1049-D1 | 0.93 | 134 | **0.351** | 0.372 | **0.277** | 0.302 |
| T1074-D1 | 0.92 | 132 | **0.427** | 0.533 | **0.269** | 0.270 |
| T1038-D1 | 0.90 | 114 | **0.503** | 0.549 | **0.290** | 0.421 |
| T1031-D1 | 0.90 | 95 | **0.639** | 0.769 | **0.547** | 0.669 |
| T1033-D1 | 0.89 | 100 | **0.513** | 0.579 | **0.673** | 0.804 |
| T1043-D1 | 0.89 | 148 | **0.569** | 0.610 | 0.469 | **0.415** |
| T1039-D1 | 0.86 | 161 | **0.619** | 0.656 | **0.466** | 0.599 |
| T1040-D1 | 0.81 | 130 | 0.653 | **0.647** | **0.552** | 0.568 |
| T1064-D1 | 0.75 | 47 | **0.491** | 0.593 | - | - |
| T1027-D1 | 0.56 | 127 | 0.626 | **0.597** | 0.523 | **0.489** |
| T1029-D1 | 0.53 | 124 | **0.697** | 0.709 | **0.510** | 0.559 |

**Table S4. Further refinement of OPUS-Rota4 based on the OPUS-RotaCM prediction for *$\chi 2$*.**

| | MAE ($\chi_1$) | MAE ($\chi_2$) | MAE ($\chi_3$) | MAE ($\chi_4$) | ACC |
|---|---|---|---|---|---|
| | | | CAMEO (60) | | |
| OPUS-RotaNN2 | 21.61 | 31.13 | 49.79 | 47.78 | 55.61% |
| OPUS-Rota4 | 21.34 | **31.13** | 49.79 | 47.78 | **57.35%** |
| OPUS-Rota4($\chi$2) | 21.34 | 32.92 | 49.79 | 47.78 | 56.71% |
| | | | CASPFM (56) | | |
| OPUS-RotaNN2 | 18.85 | 28.50 | 44.88 | 44.87 | 58.17% |
| OPUS-Rota4 | 18.46 | **28.50** | 44.88 | 44.87 | 60.42% |
| OPUS-Rota4($\chi$2) | 18.46 | 30.69 | 44.88 | 44.87 | **60.56%** |
| | | | CASP14 (15) | | |
| OPUS-RotaNN2 | 28.21 | 40.14 | 51.93 | 40.76 | 41.16% |
| OPUS-Rota4 | 28.33 | **40.14** | 51.93 | 40.76 | **43.38%** |
| OPUS-Rota4($\chi$2) | 28.33 | 41.72 | 51.93 | 40.76 | 42.69% |
